# Bacteriophage T4 gene 32 protein: Insights into its Interaction with ssDNA, binding cooperativity, and conformational change

**DOI:** 10.1101/2025.09.02.673855

**Authors:** Jules Guei, Michael P. Chapman, Paul N. Brothers, Shital Desai, Richard L. Karpel

**Author notes:** Deceased (Michael P. Chapman).

## Abstract

The single-stranded DNA binding protein of bacteriophage T4, gp32, has important roles in replication, recombination, and repair. gp32 possesses three domains: the central (core) domain which contains the binding trough for single-stranded DNA, the N-terminal domain, which interacts with the core domain of an adjacent ssDNA-bound protein, bringing about binding cooperativity, and the C-terminal domain, which interacts with other proteins involved in replication, recombination, and repair. The essential residues within the N-domain for the association with the adjacent DNA bound gp32, Lys-Arg-Lys-Ser-Thr, the *“LAST Motif”*, is almost identical to the ssDNA-interactive residues within the core domain binding trough, and was the basis of a model in which a “closed” ⇄ “open” conformational change within core domain controls DNA binding. In this study, we show that alteration of the core domain *LAST sequence*, while maintaining its *composition*, can have an effect on the binding parameters, and may be the result of a shift in the closed-open equilibrium. Additionally, utilizing a gp32 truncated at residue 227, as well as amino acid substituted variants, we have further localized the residues within the core domain responsible for the protein-protein association leading to cooperative ssDNA binding. Truncation leads to an increase in the non-cooperative affinity for single-stranded nucleic acids, which can be explained by the absence of a closed conformation in this variant. The truncated protein forms a tight complex with core domain on a 12-residue oligonucleotide, a potential candidate for further structural study.

## Introduction

Gene 32 protein (gp32), the single-stranded DNA binding protein (SSB) of bacteriophage T4, has well-established roles in DNA replication, recombination and repair, and is a model for this class of DNA-interactive proteins.[1–7] There is a large body of work delineating the details of its interaction with single-stranded nucleic acids and of the nature and function of its structural domains.[4, 5, 7–22] The protein is a single polypeptide with three domains: the basic, largely α-helical N-domain, residues 1-21, required for binding cooperativity; the globular core domain, residues 22-253, which contains the single-stranded nucleic acid binding site; and the acidic, apparently unstructured C-domain, residues 254-301, which has a major role in the heterotypic protein-protein interactions of gp32 with other T4 proteins involved in replication, recombination and repair. The three-dimensional structure of the core domain was determined by Shamoo et al.[23] (Fig 1A and Fig 1B) A large number of studies, employing various truncated forms of the protein lacking the N-, C-, or both domains clearly show that cooperative binding of the protein to single-stranded (ss)DNA involves two binding events: the aforementioned nucleic acid interaction with the core domain coupled with the association of the N-domain of an ssDNA-bound protein with the core domain of the adjacent DNA-interacting gp32.

**Fig. 1.**
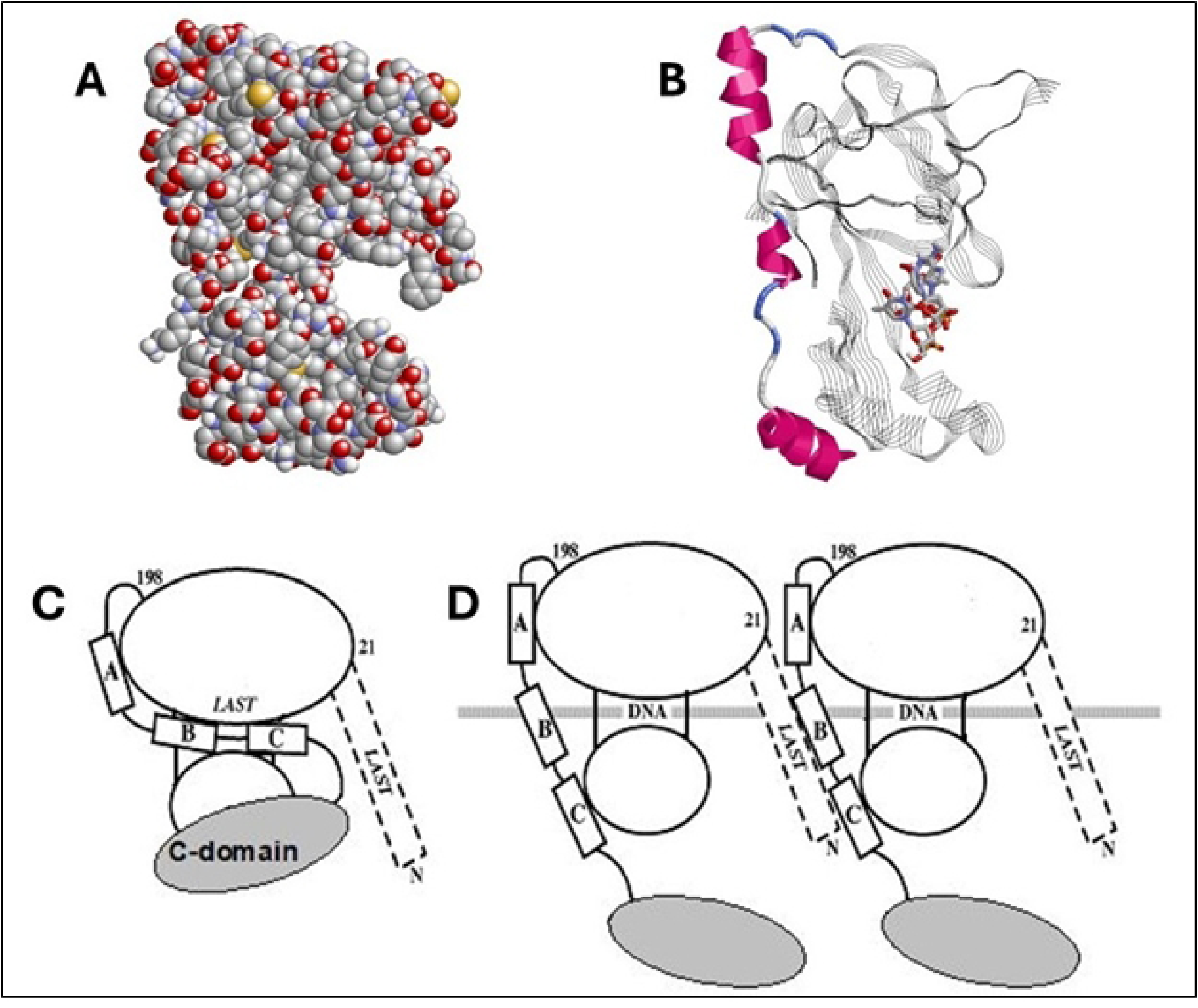
Representations of Gene 32 Protein and Core Domain structures. A. Space-filling display of core domain crystal structure [23]. This view optimizes the presumed binding trough for ssDNA, seen at the right. B. Core domain (stands and ribbons) with tetranucleotide (sticks) modeled into the trough by Shamoo *et al.* [23]. The three alpha helices shown as red ribbons correspond to residues 202-212, 216-220, and 228-237, corresponding to α-helices “A”, “B”, and “C”. C. Model for full-length gp32 in closed conformation. D. Model for full-length gp32 in open conformation, bound to ssDNA.

This apparently simple division of labor of the domains’ structure and function is somewhat more complicated: The basic amino acid sequence of the N-domain that is essential for binding cooperativity, residues 3-7 (Lys-Arg-Lys-Ser-Thr, the *“LAST’* motif[24]) is virtually identical to the sequence at the ssDNA binding trough within the core, residues 110-114 (Lys-Arg-Lys-Thr-Ser). We proposed a model in which, in the absence of DNA, an acidic region of the protein within the core domain associates with the internal (110-114) *LAST* residues, the “closed” conformation, and then upon binding ssDNA, undergoes a conformational change so that it now interacts with the N-domain *LAST* motif of the adjacent DNA-bound protein (Fig. 1C and 1D)[24]. Subsequent single-molecule DNA stretching experiments on the full-length and truncated forms of the protein provided further evidence for such a conformational change[4, 5, 25–31]. The internal protein-protein interaction unique to the closed conformation at or near the DNA binding site (internal LAST) would block binding to DNA. These observations indicate a linkage between the interaction of the protein with an isolated binding site on ssDNA, an element of both cooperative and non-cooperative binding, and the protein-protein interaction that occurs in cooperative binding. A structural change to an “open” conformation, unblocking the internal *LAST* site, is required for DNA binding. The closed conformation would also inhibit the double-stranded (ds)DNA helix-destabilizing activity of the protein, which we have shown is dependent on the (albeit weak and temporal) binding of the protein to dsDNA.[26] We note that the closed⇄open transition is salt-dependent, so that at high ionic strength, ≳0.2 M NaCl, the protein is in the open form. The model we proposed correlated with a large body of ensemble binding, helix-destabilizing, and kinetic data.[24]

In this report, we further explore details of gp32 single-stranded DNA binding by introducing variations in two critical regions of the protein. First, we determined if the exact *LAST* sequence at the core domain ssDNA binding trough, rather than its composition, is critical for ssDNA binding. Consequently, the sequence of residues 110-114, K*<u>RKT</u>*S, was scrambled to K*<u>KTR</u>*S (“KTR”) and to K*<u>TKR</u>*S (“TKR”), in each case altering the position but not the quantity of positively-charged residues. Thus, whereas in wild type gp32 the *LAST* sequence has, N to C, three consecutive positively-charged residues followed by two with hydroxyl-containing sidechains, in the two variants we now examine only two basic residues are adjacent to each other, and the – OH sidechains are now separated by one or two of the basic residues. We find that although the DNA-binding properties of the mutated proteins were generally similar to those of wild-type, the KTR variant showed a 10-fold reduction in cooperativity. This might reflect a shift toward the closed conformation in KTR, relative to the closed⇄open equilibrium in wild-type and TKR.

In a second set of experiments, the region of the core domain that interacts with the N-domain of an adjacent cooperatively-bound gp32 was probed by utilizing protein variants with amino acid residue changes and truncations in the C-end of the core domain. In this regard, we note that the classical gp32 truncate lacking the C-domain, *I (residues 1-253), maintains binding cooperativity, so the location(s) within an ssDNA-bound protein associating with the N-domain of the adjacent gp32 must be somewhere within its core domain, residues 22-253. Our new data provide further details and insight into the locations and nature of these homotypic protein-protein interactions, and we also provide interesting and correlative results on dimer formation between a *I truncate (residues 1-227) and core domain in the presence of an ssDNA 12-mer.

## Materials and Methods

### Proteins and Nucleic Acids

Gene 32 protein and the variant forms used in this study were prepared as previously described.[8, 9, 25] Mutations were introduced into plasmid pZN1, which was created by insertion of gene 32 lacking the first codon (from the expression plasmid for full-length protein, pYS6) into the Pme1 and BamH1 sites of pNEB193, a pUC19-based vector (New England Biolabs). The mutation procedure employed the QuikChange method (Stratagene), with the appropriate oligonucleotides. DNA fragments were then inserted back into pYS6. All mutations were verified by DNA sequencing (Biopolymer Laboratory at University of Maryland Baltimore). The pYS6 variants were induced, overexpressed and purified as previously described for gp32 and truncated products. [8, 9] Poly(uridylic acid), poly(U), and poly(ethenoadenylic acid), poly(εA), were obtained from P-L Biochemicals.

### Fluorescence Binding Experiments and Analyses

Binding data obtained from the quenching of protein tryptophan fluorescence by single-stranded nucleic acids were obtained by the procedure described in [32]. Aliquots of nucleic acid were added to the protein in a thermostated (20°C) SPEX Fluoromax 2 spectrofluorimeter, with excitation and emission wavelengths set at 290 nm and 342 nm, respectively. In binding experiments utilizing poly(εA)[31, 33], aliquots of protein were added to the polynucleotide solution, with excitation and emission wavelengths set at 320 nm and 420 nm, respectively. The bandwidths for both excitation and emission were 4 nm. Quartz cuvettes were used to monitor Trp fluorescence, methacrylate curvettes to monitor ethenoA fluorescence. Sample size was 1000 μL, with added aliquots generally 5 μL or less. The dependence of binding affinity on salt levels was determined by conducting binding experiments at varying [NaCl].

Fluorescence quenching and enhancement data were analyzed via the McGhee-von Hippel non-specific binding model[34]. A non-linear least squares fitting program (NFIT) was used to determine association constants. The data for K_int_ has an uncertainty of ±25% – ±33%, and for ω, ±33 – ±50%.

## Results

### Scrambling the internal LAST sequence can alter binding cooperativity

In these experiments we determined the effect of altering the *sequence*, rather than the *composition,* of the critical “*LAST*” residues within the single-stranded nucleic acid binding trough. Thus, K*<u>RKT</u>*S was altered to K*<u>KTR</u>*S (the “KTR” variant) and to K*<u>TKR</u>*S (“TKR”). Binding activity was monitored by two methods: the reduction of intrinsic protein tryptophan fluorescence upon addition of aliquots of poly(uridylic acid), poly(U), and the increase in the fluorescence of poly(ethenoadenylic acid), poly(εA), upon addition of aliquots of protein. Both these methods have been used extensively to determine binding parameters of gp32 and its classical truncated forms. At low ionic strength, high affinity conditions, 0.05 M NaCl, 0.02 M HEPES pH 7.68, 1×10^−4^ M EDTA/Na^+^, we obtained binding isotherms that initially displayed a linear decrease in Trp fluorescence (in the poly(U) titrations) or an increase in poly(εA) fluorescence up to the saturation point, followed by no further change in fluorescence. The resulting bi-linear plots yield the apparent occluded site size, with values of 8±2 nucleotides covered by each KTR or TKR protein, as is the case for wild-type protein. A typical titration, for KTR with poly(U), is shown in Fig. 2A. and for TKR with poly(εA), in Fig. 3 (insert). Although not pursued in detail, the decrease in Trp fluorescence was reversed upon addition of salt (Fig 2B), yielding a linear log K_app_/log[NaCl] plot similar to that found for gp32[16].

**Fig. 2.**
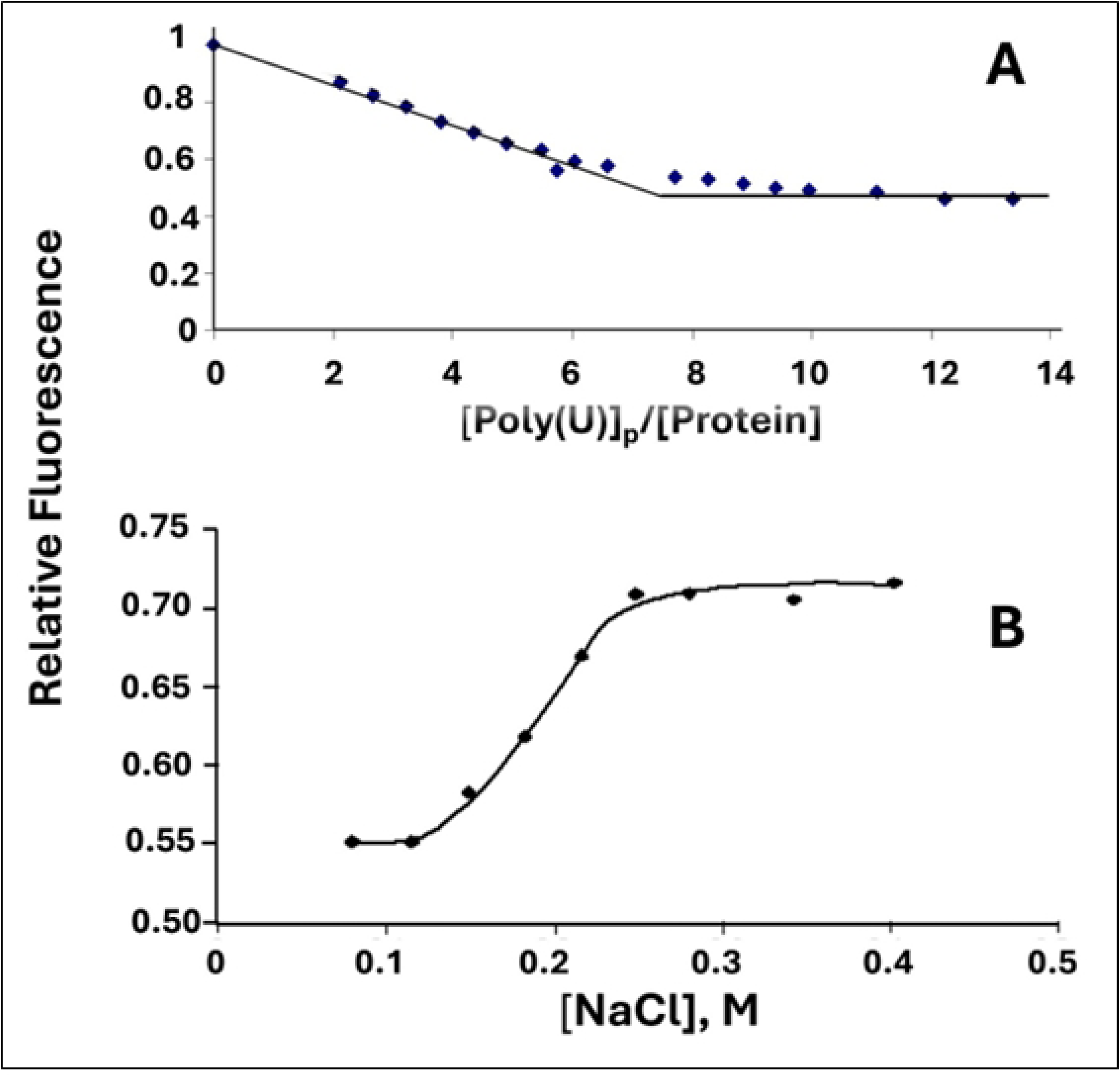
(A) Determination of the occluded binding site size of KTR at low [salt] and (B) reversal of binding by addition of NaCl. (A) [KTR] = 2.1×10^−6^ M, 0.02 M HEPES pH.7.68, 1×10^−4^ M EDTA/Na^+^, 0.05 M NaCl. (B) [KTR] = 8.6×10^−7^ M, [poly(U)]_p_ = 2.9×10^−4^ M(p), 0.02 M HEPES pH.7.68, 1×10^−4^ M EDTA/Na^+^.

**Fig. 3.**
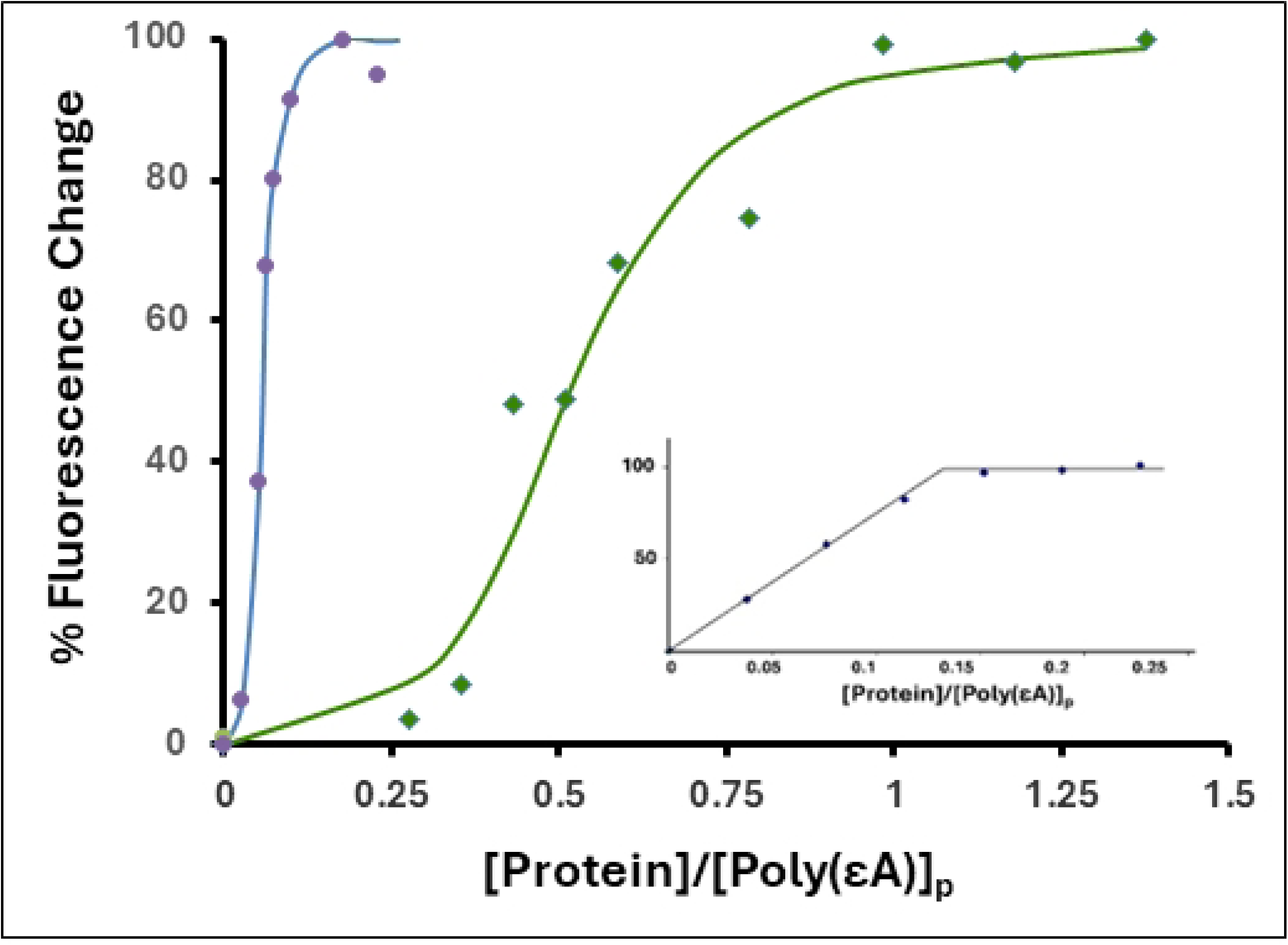
Cooperative binding of TKR (blue circle) and KTR (green diamond) to poly(εA) (representative titrations). 2.0 μM(p) poly(єA), 0.45 M NaCl, 0.02 M HEPES, pH 7.68, 1×10^−4^ M EDTA/Na^+^. *Inset:* Titration with TKR in 0.05 M NaCl.

In order to determine binding affinity, we chose to perform poly(εA) fluorescence enhancement experiments at 0.45 M NaCl, salt conditions where the affinity of protein for the polynucleotide is significantly lower than at 0.05 M NaCl. The binding isotherm obtained at 0.45 M NaCl with gp32 is clearly sigmoidal, indicative of cooperativity [14]. The cooperativity parameter, ω, the probability of a protein binding next to an already-bound protein relative to binding anywhere on the single-stranded nucleic acid, has a value of ∼1000 for gp32, and is independent of base composition, sugar (ribo or deoxy), or [salt][14, 16]. Addition of aliquots of KTR or TKR to poly(εA) produced sigmoidal binding plots qualitatively similar to those seen with gp32. Representative titrations are shown in Fig. 3.

The binding data was analyzed via the cooperative binding model of McGhee and von Hippel (eq. 15)[34], leading to calculation of K_int_, the association constant of protein for an isolated binding site on nucleic acid, and ω (Table 1). The cooperativity associated with KTR binding is about an order of magnitude lower than the value for TKR or gp32, and the overall binding affinity for KTR, K_int_ω, is about 4-fold lower. These trends were seen when the data was analyzed with a range of occluded size sizes, n, which is generally considered to be 8±1 nucleotides. Note that at 0.45 M NaCl, the literature value of K_int_ω for unmodified gp32 binding to poly(εA) is 7.0×10^6^ M^−1^, within the uncertainties of measurement comparable to the TKR result.[14]

**Table 1.** Binding parameters of the gp32 variants TKR and KTR. Calculated with occluded binding site size, n = 8. 0.45 M NaCl, 0.02 M HEPES pH 7.68, 1×10^−4^ M EDTA/Na^+^. Uncertainties for K_int_ were 25% (TKR) and 33% (KTR), for ω were 50% (TKR) and 33% (KTR), and for K_int_ω were 50% and 33% respectively.

| | $K_{int}, \text{M}^{-1}$ | $\omega$ | $K_{int}\omega, \text{M}^{-1}$ |
| --- | --- | --- | --- |
| **TKR** | **$2.2 \times 10^3$** | **1500** | **$3.6 \times 10^6$** |
| **KTR** | **$6.5 \times 10^3$** | **160** | **$9.3 \times 10^5$** |

As we have noted, under the salt conditions of these experiments (0.45 M NaCl), gp32 is in the “open” conformation, so that the energetics of the intermolecular protein-protein binding event, leading to cooperativity, is independent of the need to break up the intramolecular binding of the “closed” conformation. The reduced value of ω and slightly higher value of K_int_ seen for KTR, relative to that of TKR and gp32 is thus somewhat problematic, since the cooperativity of binding, the result of protein-protein interaction, would seem to be independent of the (altered) *internal LAST* sequence involved in the association with ssDNA (the N-terminal *LAST* sequence, KRKST, participating in the homotypic protein-protein interaction, was not changed in either of the variants). Conceivably, the altered internal *LAST* sequence in the KTR variant possesses a stronger binding affinity in its interaction with an *intra*molecular acidic region. This would shift the equilibrium toward the proposed closed conformation so that the protein population is not totally in the open form, therefore reducing the magnitude of ω. At the same time, the increased affinity for ssDNA would compensate partially for the increased competition by protein intramolecular binding.

### Cooperativity: The region of the core domain associating with the N-domain of an adjacent DNA-bound protein is likely within residues 198 – 253

The segment of core domain that is in contact with the N-domain of an adjacent protein cooperatively-bound to ssDNA is unknown, but, given the basicity of the N-domain *LAST* residues, this region is likely to be acidic. In this regard, the sequence from Pro-198 to the C-terminus of the core (Lys-253) is acidic, with 9 (Asp + Glu) and 5 Lys residues, and consists of three α-helical segments, residues 202-212, 216-220, and 228-237, respectively labeled “A”, “B”, and “C”, and connected by shorter, flexible segments (Fig. 1). This region is largely located at the exterior of the protein, and therefore constitutes a potential candidate for the surface which binds the adjacent protein’s N-domain. Interestingly, it is immunologically cross-reactive with ssDNA [18]. Thus, its participation in protein-protein interactions would mimic the binding of core domain *LAST* residues to ssDNA (Fig. 1, C and D).

Two approaches were used to localize this interactive surface. In the first case, via insertion of stop codons following those coding for residues 201, 216, and 227 in full-length gp32, deletion mutants were constructed with termini corresponding to elimination of three, two, and one of the α-helices, respectively. High levels of expression were obtained, but we were able to solubilize only the protein corresponding to loss of the helix closest to the C-terminus, the “C” helix in Fig. 1, 1-227. In the second approach, two gp32 variants were constructed with elimination of the negative charge at positions within and adjacent to the second (“B”) helix (D215A, E218A, D223A) and at positions within “C”, the helix closest to the C-terminus (E229A, E230A). In this case, both of the expressed proteins were fully soluble.

To determine the effect of the truncation and substitutions on binding cooperativity, we observed the association of the proteins to poly(ethenoadenylic acid), poly(εA). As we have seen with gp32, KTR, and TKR at 0.45 M NaCl, upon addition of protein the increase in polynucleotide fluorescence displays the classical sigmoidal shape associated with highly cooperative binding [14]. However, when the 1-227 truncate was added to poly(εA) at 0.45 M NaCl, the binding was no longer sigmoidal, and analysis of the isotherm indicated a loss of all, or most of the cooperativity (ω ≈ 10). (Fig. 4, Table 2) This indicates that homotypic protein-protein interaction requires an intact C α-helix, residues 228-237, and possibly residues between this helix and the C-terminus at residue 253. Curiously, the E229A, E230A variant, with its loss of the two negatively charged amino acids within this helix, did not lose cooperativity. A somewhat different result was obtained with the D215A, E218A, D223A variant, corresponding to the loss of three negative charges adjacent to and within the 216-220 B (middle) helix. In this case, the binding isotherm is still clearly sigmoidal, though there is a moderate reduction in cooperativity (ω ≈ 100).

**Fig. 4.**
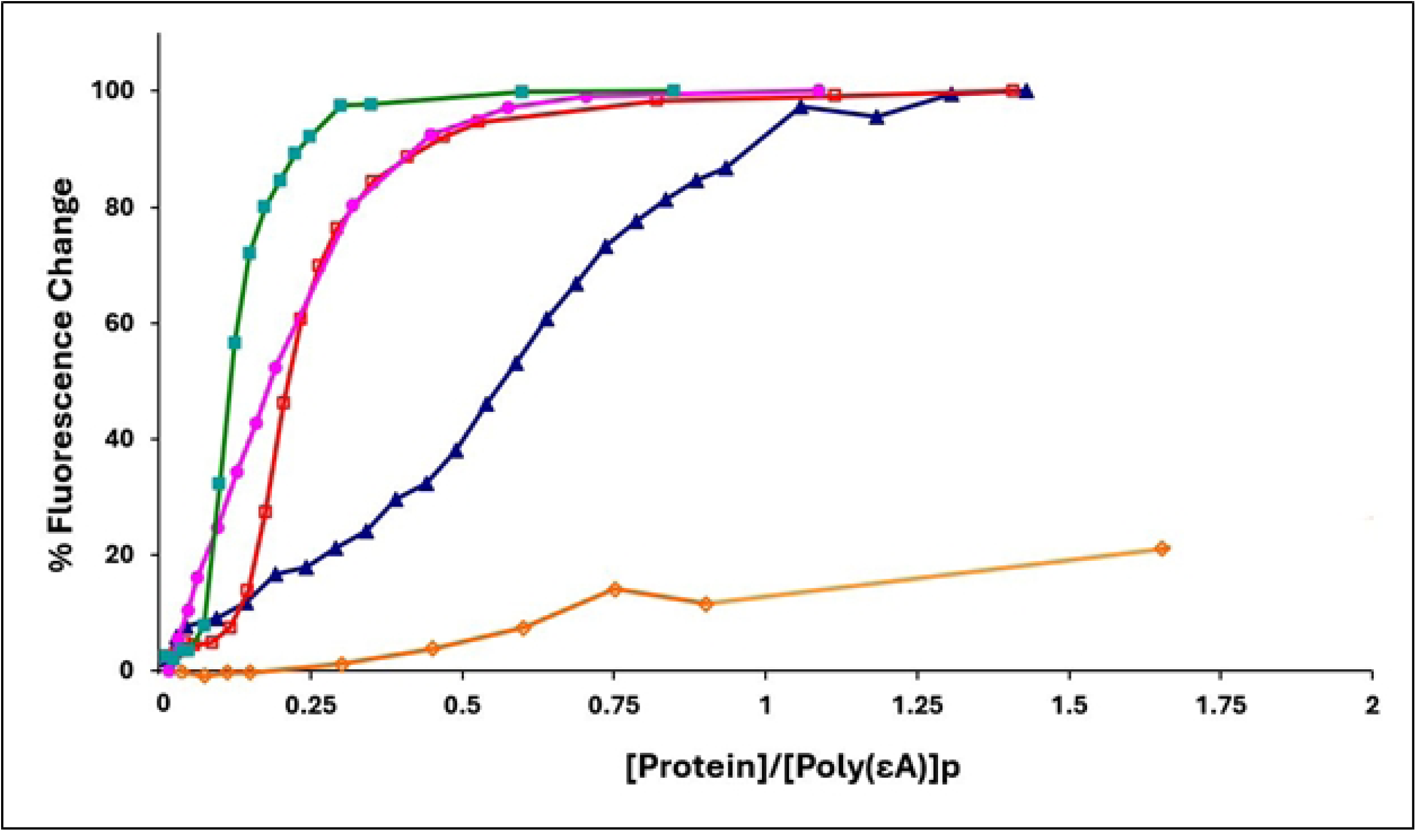
Binding of gene 32 protein variants to poly(εA) (representative titrations). 0.45M NaCl, 0.02 M HEPES, pH 7.68, 1×10^−4^ M EDTA/Na^+^. Green square, gp32; pink circle, 1-227 truncate; open red square, E229A, E230A; blue triangle, D215A, E218A, D223A; open orange diamond, *III (core domain)

**Table 2.**
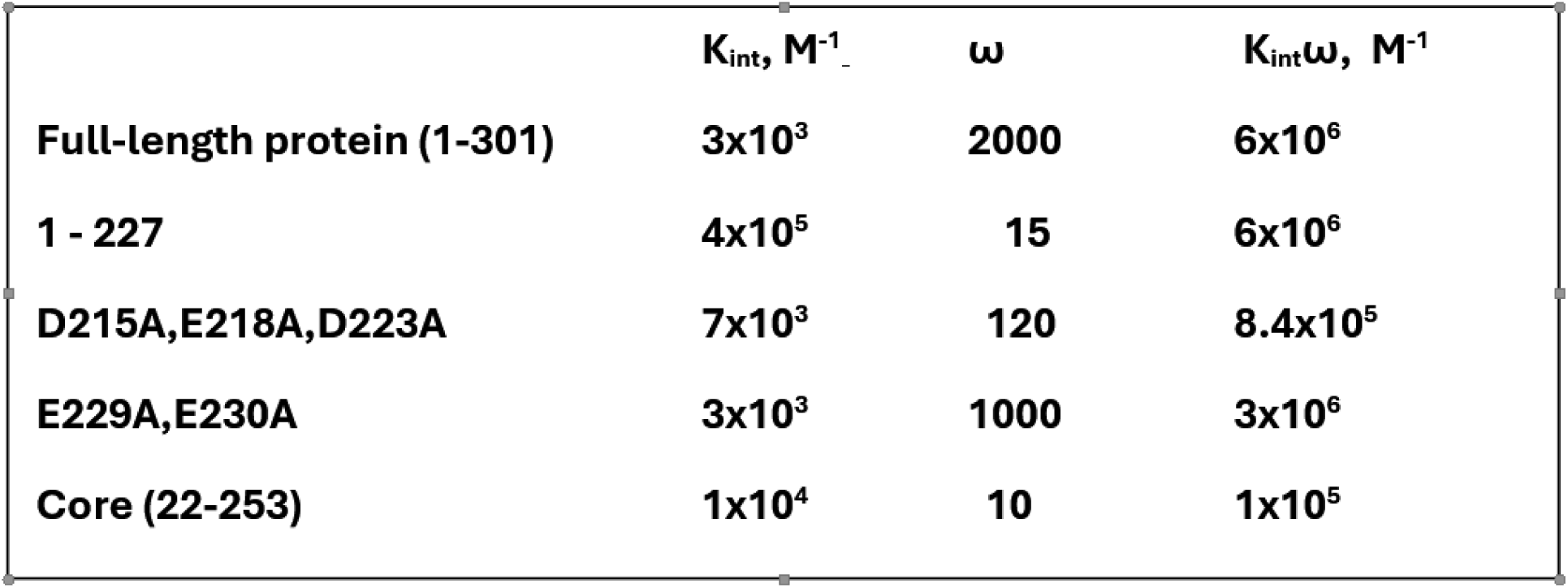
Binding constants of gene 32 variants with poly(εA) in 0.45 M.

These results add to our understanding of the nature of the homotypic protein-protein interaction that leads to binding cooperativity. The simplest interpretation of the loss of cooperativity in the 1-227 variant is that the surface of core domain interacting with the N-domain of an adjacent-bound gp32 is located between residues 228 and 253 (the C-terminus of the classical *I truncate which retains full cooperativity).

Alternatively, loss of this section of might alter the structure of other locations within the protein that are required for protein-protein association. However, in the experiments described below, we show that 1-227 still retains its ability to associate, via its N-domain, with intact core domain (*III) while binding to a single-stranded oligonucleotide, so any major alteration of structure is highly unlikely.

### NaCl

As we see in Table 2, the 1-227 variant shows a striking two-fold order of magnitude increase in K_int_ relative to the full-length protein, as well as to the other proteins examined. In the model shown in Fig.1C and 1D, the binding of gp32 to ssDNA requires the removal of the internal interaction involving the ssDNA binding trough *LAST* residues. This presumably requires an input of energy to break the internal attraction, which would reduce the overall free energy of binding to an isolated site on ssDNA (-RTlnK_int_). Unlike gp32 and all the other variants examined, 1-227, truncated between the B and C helix, might not undergo this interaction, would then not need the associated input of energy, and therefore would possess a larger affinity for single-stranded nucleic acid.

The removal of negative charge within or adjacent to the B helix, residues 216-220, and the C helix, 228-237, leads to several possibilities. Given the essential participation of the “*LAST*” residues of the N-terminus, KRKST, we had expected that electrostatic interactions with acidic solvent-exposed residues at these locations were critical for cooperativity. The simplest explanation for the lack of any effect on the cooperativity parameter, ω, for 228-237, and only a small reduction for 216-220 is that electrostatic interactions are not involved. This is consistent with the observed absence of a salt dependence of ω [14]. Moreover, T4 phage with Arg→Ser or Arg→Tyr substitutions at residue 4 of gp32 (within the N-domain *LAST* sequence) were viable, suggesting that hydrogen binding ability is more critical than or at least substitutable for charge at this position (Karpel *et al*, unpublished results).

There are other possible explanations for our results. Referring to our model [24], the electrostatic interactions lost upon dissociating the *intra*molecular interaction (Fig. 1C) would be compensated by the *inter*molecular interactions acquired upon binding ssDNA cooperatively (Fig. 1D). There would be no net change in ionic bonds, and ω would therefore be salt-independent. Whatever the nature of the weak chemical interactions between the N-domain and core, we conclude that the segment of core between residues 227 and 253 is essential for bringing about binding cooperativity.

The linear salt dependence of K_int_ for the binding of poly(εA) to the two substitution variants, E229A,E230A and D215A,E218A,D223A, is essentially the same as that of gp32 (Fig. 5). The slope of the logK_int_ vs. log[NaCl] plot for these three proteins is –3.2. The 1-227 variant, which has largely lost both its ability to bind cooperatively and, as we propose, the internal *LAST* interaction, shows a consistently larger K_int_ over the range of [NaCl], 0.05 to 0.45 M NaCl. The slope of the plot for 1-227, –2.4, is somewhat smaller than that of the other proteins. Record and co-workers have shown that the log-log dependence of affinity on salt concentration is a function of the number of ion-pairing interactions between nucleic acid and protein (related to the release of cations from the nucleic acid upon binding) and the number of cations displaced from protein.[35] It is difficult to predict how the conformational change we propose for gp32 (Fig. 1, panels C and D) would affect the net number of cations and anions released, but it is noteworthy that the two substitution variants behave comparably to unmodified protein. The different slope seen with the truncated 1-227 might reflect the loss of the internal *LAST* interaction.

**Fig. 5.**
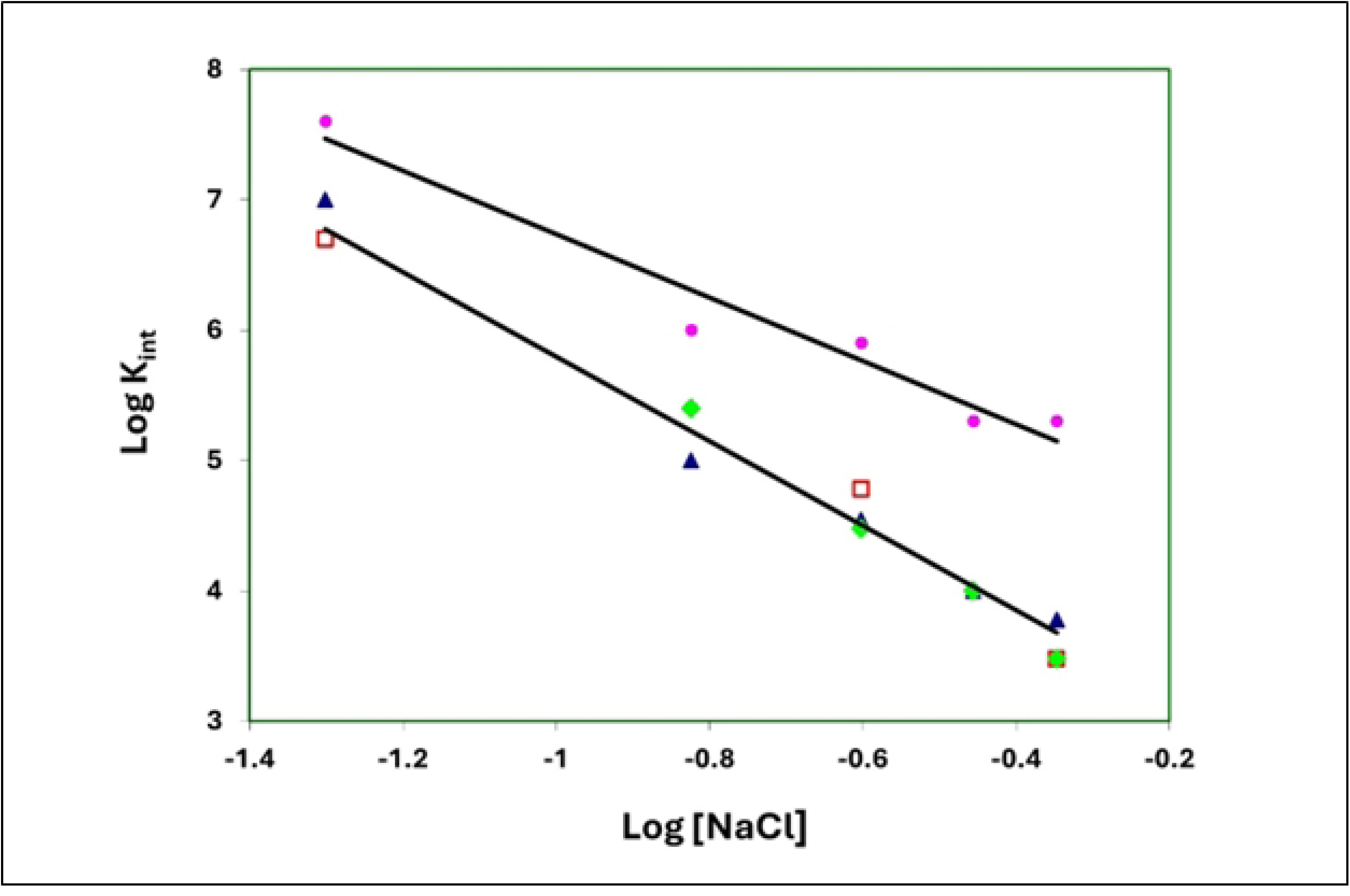
Binding to poly(εA): Salt dependence of K_int_. Open red square, gp32; blue triangle, 215A,E218A,D223A; green diamond, E229A,E230A; pink circle,1-227. At 0.25 and 0.35 M NaCl, E229A,E230A and D215A,E218A,D223A yield essentially the same K_int_, and at 0.45 M NaCl, the K_int_ values for gp32 and E229A,E230A are also nearly identical.

### Core domain promotes the binding of the 1-227 truncate on a 12-residue oligonucleotide

Neither the *III (core domain) nor the 1-227 truncate are capable of binding single-stranded nucleic acids cooperatively, the former lacking the N-domain, the latter lacking the portion of the core needed for protein-protein interaction. However, the N-domain is present in 1-227, and the interactive surface is intact within *III. Therefore, 1-227 should be capable of binding to *III while both are bound to single-strand DNA or RNA (Fig. 6). In order to determine if this is the case, we explored the binding of 1-227 and *III to a 12-residue oligodeoxynucleotide capable of binding two protein molecules, d(εA)_11_T[14]. As with the poly(εA) polynucleotide, binding is followed by the enhancement of ethenoadenylate fluorescence.

**Fig. 6.**
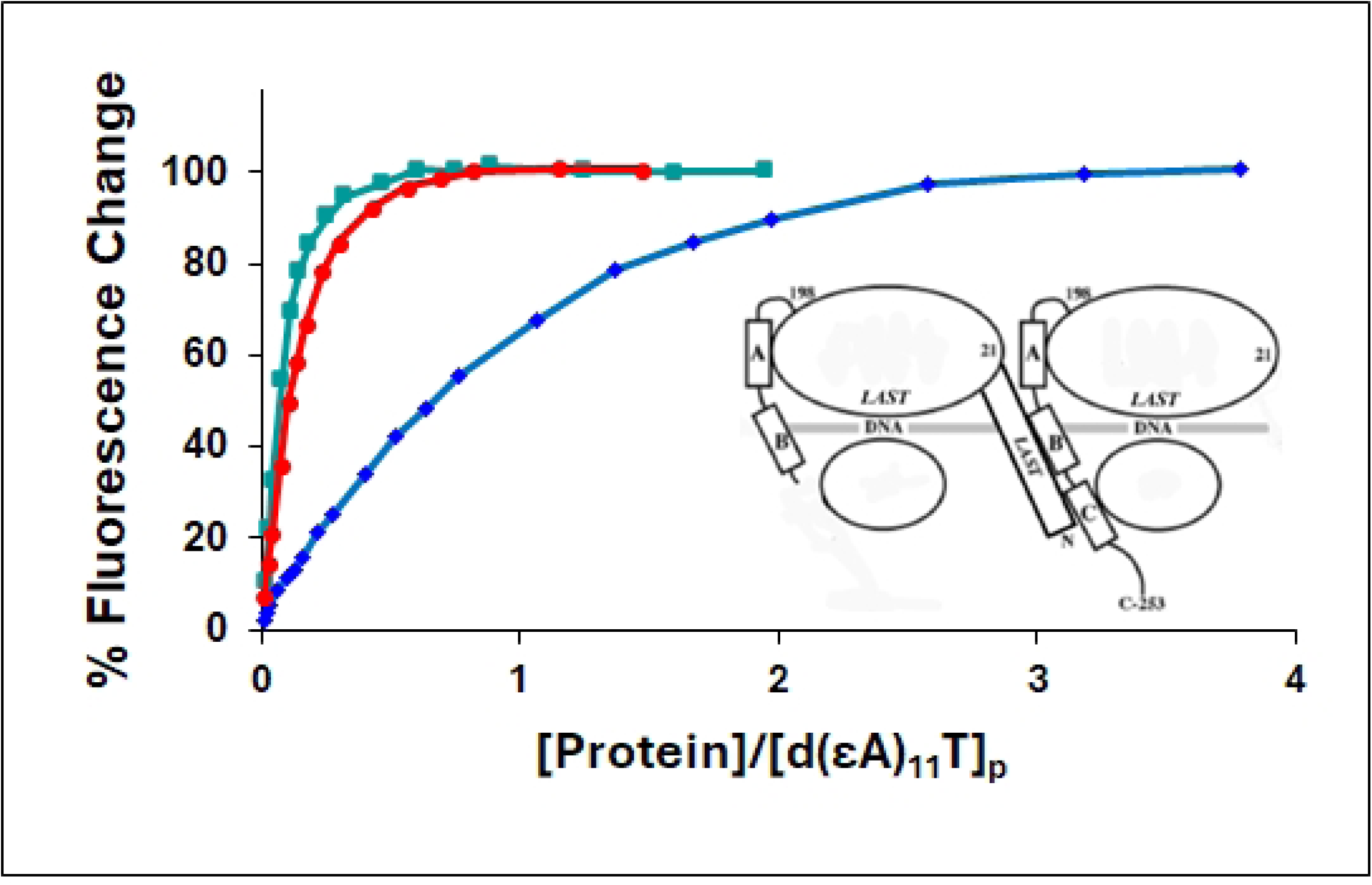
Recruitment of 1-227 by *III (core domain). Binding of proteins to d(εA)_11_T in 0.01 M NaCl, 0.02 M HEPES, pH 7.68, 1×10^−4^ M EDTA/Na^+^, monitored by the increase in oligonucleotide fluorescence. Green square, *III; blue diamond, 1-227; red circle, *III + 1-227 (1:1). For the *III + 1-227 points, the concentration of protein was the sum of the *III and 1-227 values, i.e., total protein concentration. Inset: Left: 1-227; Right: *III.

Under low salt conditions, 0.01 M NaCl, *III on its own binds strongly to d(εA)_11_T, whereas 1-227 demonstrates relatively weak binding (Fig. 6). A striking result was obtained when a 1:1 mixture of *III and 1;227 was added to the oligonucleotide. In this case, the oligonucleotide fluorescence increase is well in excess of what would occur from the sum of the individual changes effected by *III and 1-227, leading to a binding isotherm nearly identical to that of *III. Thus, the presence of core domain enhances the binding of 1-227. The synergistic effect is very likely the result of a cooperative interaction between 1-227 and *III, the N-domain of the former in contact with a binding surface located between residues 227 and 253 of the latter (and possibly other sites within the core domain). A very similar result was obtained at 0.05 M NaCl (not shown).

We note that while 1-227 had a greater affinity for d(εA)_11_T than did *III under conditions of low salt, the order was reversed for the binding to poly(εA) at 0.45 M NaCl. Obviously, there are major differences in ionic strength, length of the nucleic acid, and the *deoxy* oligo vs. the *ribo* polymer. Each of these factors could influence affinity, but they do not affect our interpretation of the results shown in Fig. 6.

## Discussion

In this study, we have explored the involvement of two regions of gene 32 protein that are critical for its varied binding activities. These are (a) the ssDNA binding trough within the protein’s core domain, and, (b) the surface of the core domain that is in contact with the N-domain of an adjacent ssDNA-bound protein, which brings about cooperative binding.

With respect to (a), the critical amino acid residues within the ssDNA binding trough are residues 110-114, Lys-Arg-Lys-Thr-Ser, the *LAST* Motif [24]. In this report, we show that scrambling this sequence can have a small effect on the affinity of the protein for an isolated binding site, K_int_, on single-stranded nucleic acid, and a somewhat larger effect on cooperativity. This is a result consistent with the biological activities of gp32, where the protein encounters a wide variety of sequences within its ssDNA substate. During DNA replication, for example, gp32 traverses single-stranded DNA along the entire length of the bacteriophage T4 genome, so the placement of positive charges and hydrophilic sidechains capable of hydrogen bonding optimized for one particular ssDNA sequence would not necessarily be optimized for a different sequence. For each placement, the average of affinities would likely be similar.

We observed a ∼10-fold lower cooperativity parameter for the KTR mutant relative to that of TKR and gp32. As we have noted, within the ssDNA binding trough, of the three basic residues within the “*LAST*” sequence, two are adjacent in KTR and TKR, three in gp32. The lower cooperativity of KTR, reflecting intermolecular protein-protein association, cannot be attributed to this basic residue placement. In contrast, the arrangement of positive charges within the ssDNA binding trough *LAST* sequence might be critical for the proposed internal protein-protein closed interaction, moderately stronger in the case of KTR. Within the trough, the position of the ssDNA is not rigid, as evidenced by the structural disorder observed crystallographically [23], and, as we have noted, functionally necessitated by the protein’s need to interact with the entire T4 genome (as ssDNA). However, this would not be the case for the internal protein-protein binding of the closed conformation, where there would likely be a specific set of ion-pairing, hydrogen bonding, and/or non-ionic interactions.

We note that, in contrast to the arrangement of positive charges within the protein’s binding trough, the location of negative charges within the ssDNA substrate directly interacting with protein is very significant. We showed that at least two, and likely three adjacent phosphodiester linkages are a minimal requirement for binding single-stranded nucleic acids.[9] This observation is in concert with the finding that two to three nucleotides are directly involved in the binding interaction.[19] Utilizing 2-aminopurine-labeled substrates, von Hippel and co-workers have established the orientation of single-stranded DNA substrates within the binding groove, with the 5’-end closest to the N-domain/core domain juncture, and the 3’-end closest to the C-domain/core juncture.[10, 12, 19]

In addition to electrostatic interactions, protein aromatic residues likely also interact with ssDNA bases[10, 23, 36], and non-charged residues could play a role in stabilizing the closed conformation. This is clearly also the case for the intermolecular protein-protein interaction responsible for binding cooperativity, where the positively-charged N-domain binds to a region in the adjacent DNA-bound protein that is likely between residues 227 and 253, near the C-terminus of the core domain (point (b)). The loss of two negative charges within the “C” helix, residues 228-237, has essentially no effect on cooperativity, which could indicate that uncharged residues play an important role in the intermolecular protein-protein binding. The C-domain itself is highly negatively-charged, but it is not involved in the homotypic gp32-gp32 interaction, as its loss does not remove or reduce cooperativity.[16]

Our data on the binding of the 1-227 variant strongly implicate the region between residue 228 and 253 as the location for the interaction with the N-domain of an adjacent ssDNA-bound protein, which brings about binding cooperativity. 1-227 still retains its ability to interact with that location, which is dramatically demonstrated by its recruitment by core domain, *III, to bind tightly to the 12-residue oligonucleotide, d(εA)_11_T. This complex, with the ssDNA binding functionalities of its two protein components preserved, would be a good candidate for a structural study aimed at determining the fine details of the homotypic protein-protein interaction responsible for binding cooperativity.

Gene 32 protein, the classical single-stranded DNA binding protein, has been the subject of study for more than half a century, producing a rich body of knowledge relevant to our understanding of DNA metabolism. This report furthers our understanding of this most important protein. Nevertheless, important details about its various interactions with DNA, and the concomitant effects on its biological roles, remain to be explored. In this regard, recent single molecule optical tweezer and atomic force microscopy methods have explored gp32-filament formation, higher-order structure not previously observed via ensemble studies.[37, 38] The results have included the observation that high gp32-ssDNA saturation leads to filament unwinding and rapid removal of the protein from the complex, a potentially useful property in that it would facilitate dissociation of gp32 in front of DNA polymerase during replication.[37] This was not observed with *I, the truncate lacking the C-domain, suggesting that this part of the full-length protein destabilizes the filament at high saturation levels.[38] Clearly, there are many details, yet to be discovered, of gene 32 protein, its domains and conformational changes, that are relevant to the protein’s various biological functions.

## Acknowledgements

Funding for this project was provided by NIH (GM 52049, R.L.K), and the UMBC Designated Research Initiative Fund (R.L.K.). The funders had no role in the design, data collection and analysis, decision to publish, or preparation of the manuscript. Michael P. Chapman passed away before submission of this manuscript. Richard L. Karpel accepts responsibility for the integrity and validity of the data collected and analyzed.

## Author Contributions

JG, MPC, PB, and RLK designed the study; JG, MPC, PB, and SD carried out the experiments, JG, MPC, and RLK wrote portions of the manuscript, and RLK edited and produced the final manuscript.

## Data Availability

All relevant data are within the paper.

